# Patch-DCA: Improved Protein Interface Prediction by utilizing Structural Information and Clustering DCA scores

**DOI:** 10.1101/656074

**Authors:** Amir Vajdi, Kourosh Zarringhalam, Nurit Haspel

**Affiliations:** Computer Science Department, University of Massachusetts Boston, Boston, 02125, USA; Mathematics Department, University of Massachusetts Boston, Boston, 02125, USA

## Abstract

Over the past decade there have been impressive advances in determining the 3D structures of protein complexes. However, there are still many complexes with unknown structures, even when the structures of the individual proteins are known. The advent of protein sequence information provides an opportunity to leverage evolutionary information to enhance the accuracy of protein-protein interface prediction. To this end, several statistical and machine learning methods have been proposed. In particular, direct coupling analysis has recently emerged as a promising approach for identification of protein contact maps from sequential information. However, the ability of these methods to detect **protein-protein inter-residue contacts** remains relatively limited.

In this work, we propose a method to integrate sequential and co-evolution information with structural and functional information to increase the performance of protein-protein interface prediction. Further, we present a post-processing clustering method that improves the average relative F1 score by 70 % and 24 % and the precision by 80 % and 36 % in comparison with two state-of-the-art methods PSICOV and GREMLIN.

## Introduction

Proteins interact with one another as part of their function [15]. While computational and experimental approaches for identifying the 3D structures of protein complexes has advanced significantly, there are still many complexes with unknown structures, even when the structures of the individual proteins are known. Over the past few years, there has been a growing interest in methods that apply evolutionary information extracted form protein sequence changes to protein structural and functional problems such as protein folding and protein-protein interactions.

### Existing work

Interface prediction methods can be broadly divided into template-based (structure) predictors, sequence-based methods, and hybrid methods. When structural information is available, it can be used to improve the protein-protein binding prediction. For example 3D-based predictors rely on the assumption that the binding interface has distinct structural properties compared to the rest of the protein [1, 28, 27, 33]. These models use the 3D structure of two proteins to find an optimum docking model. The main disadvantage of these methods is that they require a high degree of similarity between template structures and the target complex. Also, these models are designed based on a specific knowledge of the physical system, which is error-prone and does not generalize well.

The second category are sequence-based predictors. These predictors utilize machine learning and probabilistic models such as Bayesian methods, Hidden Markov Models (HMMs), and conditional Random Fields [23, 19, 2, 6], trained on features extracted from the proteins’ sequences to predict the binding sites. For example, sequential features such as hydrophobicity, which plays an important role in stabilizing protein-protein interactions, and amino acid propensity, are typically utilized to infer binding site properties. These features reduce the search space to specific regions in the protein and limit the putative binding sites to a smaller set of feasible residues that have interface properties. While protein interfaces are usually associated with these properties, they are not universally applicable to all complexes and associations may vary based on interacting protein families.

The third category of methods combine the previous two approaches in predicting the binding residues [9, 28, 24]. Although these methods have a higher performance rate in cross-validation studies, their performance decreased significantly when applied to new independent protein families, presumably due to high number of parameters and over-fitting.

In [5], Esmaielbeiki and others provide a comprehensive benchmark, comparing more than 70 protein interface prediction methods across multiple datasets. The conclusion of this extensive study is that the aforementioned structural features are necessary but not sufficient in determining protein-protein interactions. These features are too general and in order to have a robust and generalizable model, the characteristics of the partnering proteins should be taken into consideration for predicting the binding interface. Studies show that the most informative structural and sequential features for protein interfaces are the solvent accessible surface area (SASA), secondary structure, the geometric shape of the protein surface, hydrophobicity, conservation, amino acid propensity and the crystallographic B-factor [7, 5]. Other features have also been considered but improvement in performance is negligible.

### Protein Complexes and Co-Evolution

Protein complexes can be divided into two categories: Obligatory and Transient. All homodimer and some heterodimer complexes are examples of Obligatory complexes. Transient protein complexes form and break down transiently *in vivo*. An interaction between an enzyme and an inhibitor is an example of a Transient complex. In protein-protein interaction context the term “hotspot” refers to a residue that makes a major contribution to the binding free energy [36]. Hotspot residues help in stabilizing the protein during conformational changes [16, 32]. Protein interfaces usually appear as 3-dimensional patch of nearby residues with every patch containing a hotspot residue. The hotspot residues are typically highly conserved and the neighboring residues usually co-evolve with the residues of the interacting protein partner. It is generally believed that when two proteins interact, some of their amino acids tend to *co-evolve*, especially in Obligatory complexes since the rate of mutation in obligatory complexes is higher than Transient complexes [16, 21]. Since sequence information is much more readily available than structural information, many methods attempt to estimate the interactions between proteins using multiple sequence alignment (MSA). Existing approaches use various methods of statistical analysis to detect signals among co-evolving residues. These methods utilize Mutual Information, phylogenetics, undirected graphical models, deep learning, and more [10, 13, 4, 26, 26]. The idea of identifying co-evolving residues from MSAs has attracted special attention in recent years. Some of these methods include pseudo-likelihood-based (Markov Random Field) methods such as GREMLIN, EV-Complex, CCMPred, plmDCA or Gaussian Graphical Model (GGM) methods such as PSICOV, mfDCA [4, 29, 12, 22, 13]. These methods are designed for identifying protein contact maps (folding structure).

### Graphical Models

In recent years, multiple groups have investigated the integration of prior knowledge into both pseudo-likelihood and GGM methods for protein contact prediction, either through using machine learning approaches such as deep learning and random forest or as a penalized optimization indirectly. For example, PconsC combines PSICOV and plmDCA into a random forest as well as a 5-layer neural network in the latest release[30]. Meta-Psicov utilizes two stages of the neural network to improve upon PSICOV. In the first stage, PSICOV, mfDCA, and CCMPred are combined and utilized to predict contact sites. Later, in the second stage, the predictions are refined based on other features such as amino acid propensity, hydrogen bonds, and secondary structure [14]. CoinDCA [20] incorporates prior knowledge indirectly as a separate step into the graphical model. This model is based on PSICOV and incorporates a group-lasso penalty in the optimization function. It additionally utilizes supervised priors from protein propensity, sequence profile, mutual information, and other features to improve the predictions.

Recently, some groups have explored deep learning methods for predicting proteins’ structure and interface. RaptorX has been one of the state-of-the-art methods for predicting protein contacts [34]. It combines threading and machine learning to improve the alignment quality. Later, the same group applied convolutional neural network along with CCMPred, which resulted in improved model performance [34].

All these methods depend heavily on the quality of the MSA. Note that if a protein is highly conserved across multiple families, there is no signal for co-evolution. As a result all of these methods have MSA and phylogenetics dependency biases among different species. Moreover, indirect couplings that is typically captured as covariation, may mislead the prediction. Su lkowska et al [31] showed that not all the covariations that can be measured by the MSA correspond to co-evolution, which is one of the contributing factors behind the high false positive rates of these methods. Consequently, careful integration of a prior interaction map, learned from other non-MSA based method or clustering of top co-evolving residues can significantly improve the performance of such models.

### This Contribution

In this work, we introduce a meta-method based on undirected graphical model to improve protein contact prediction. Our focus is to directly integrate prior information into graphical models to predict the binding site between two interacting proteins. Figure 1 depicts the workflow of our proposed method. Our method directly integrates the structural information of the two interacting proteins along with docking pattern as prior knowledge into the prediction task. More specifically, the additional structural information is formulated as a penalty that is passed into a Gaussian Graphical Model (GGM). Further, we introduce a post processing step that takes the top co-evolving pairs of residues between the two proteins and constructs interface patches. While the idea of augmenting learning algorithms with auxiliary information and additional features to learn protein contacts has existed before, the integration of auxiliary information is typically **indirect to the DCA models** [35, 14, 20]. In our approach, we integrate additional sources of information, which are generalizable to all other protein families, and prior knowledge directly during the learning stage and not as a separate post processing stage. This integration can be expanded to include different sources as well.

**Figure 1.**
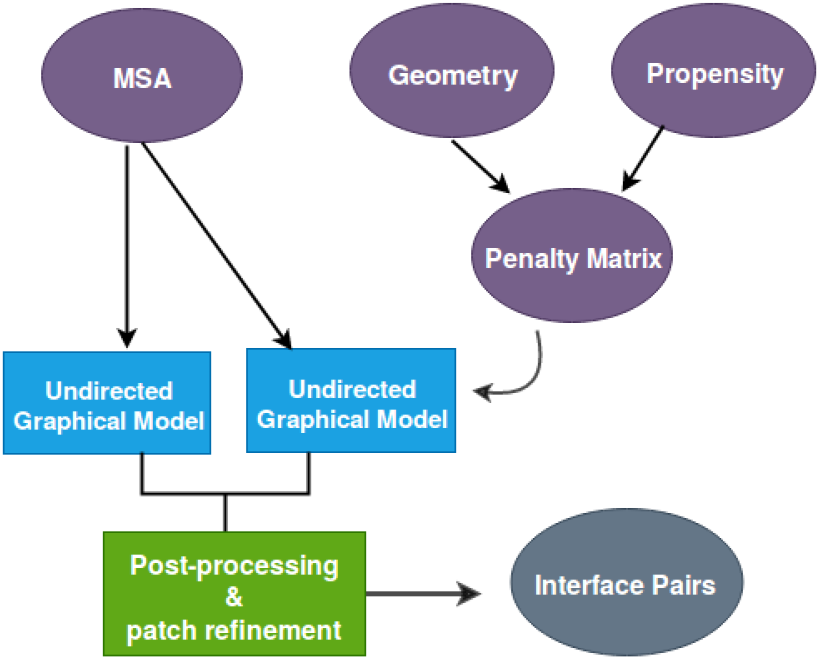
Integration of structural information into co-evolution based graphical models and patch building to filter false positive pairs among top co-evolving residues.

## Materials and Methods

We start by giving a general overview of our approach. Given two interacting proteins *A* and *B*, let *L*_*A*_ and *L*_*B*_ denote the lengths of proteins. we first construct a separate MSA for each protein using homologous sequences. We then concatenate the MSAs such that every homologue sequence of protein *A* from species *s*_1_ is matched with its corresponding homologue sequence in protein *B* in the the same species. A filtering process is applied to remove the “duplicate” sequences that are 90% or more identical to other sequences in the MSA. Additionally, columns in the MSA that have more than 75% gaps are removed. This filtering process greatly enhances the co-evolution signal in the MSA. Let *n*_*M*_ denote the number of sequences in the MSA after filtering. The concatenated MSA is then represented as a binary matrix *X*_*AB*_, where every position (amino acid) in the MSA is mapped into a binary vector of size 21 (20 amino acids and one gap), resulting in a matrix with *n*_*M*_ rows and (21 × (*L*_*A*_ + *L*_*B*_) columns. The goal of the GGM is to estimate the covariance matrix Σ and further calculate the precision matrix Θ = (Σ)^*−*1^, using which the interactions between the two proteins are inferred. Every element of the matrix Σ gives the covariance of amino acid type *a* at position *i* with amino acid type *b* at position *j*. Therefore, If the (*i, j*)^*th*^ element of Θ is zero, then positions *i* and *j* are conditionally independent, given the other positions of the MSA. Equation 1 gives the objective function of the GGM where *S* = *COV* (*X*_*AB*_), Θ, and Λ are the empirical covariance, precision matrix, and the penalty matrices respectively. The matrix Λ controls the sparsity of the precision matrix. High values of Λ shrink the corresponding value of the precision matrix to be 0.

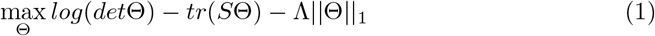

In our method, the matrix Λ is constructed based on the structural features of the two contacting proteins using several sources of information that are combined together by a weighted ensemble average. One of the advantages of constructing a matrix Λ in this fashion is that multiple sources of information can be combined and integrated into the optimization. In this work, we use two sources of information. Other sources can be added in the same fashion.

### Integrating geometry-based information

Since much of the interaction between pairs of proteins is determined by geometric complementarity of their surfaces, we incorporated geometry based scores into the penalty matrix. We utilized the ClusPro docking algorithm [18] to calculate geometric complementarity models of the two proteins. The ClusPro algorithm performs the following steps:

1. Fast Fourier Transform (FFT) based search [17]. One protein is placed on a fixed grid and the other on a movable grid, and the search is conducted based on geometric and energetic constraints.
2. Clustering the resulting conformations based on Interface RMSD (IRMSD).
3. Filtering and refinement to remove steric clashes.

The algorithm returns the top-*k* complexes, where *k* is a user-specified value and is set to 10 by default. We initially used *k* values ranging from 50 to 150. The top candidate complexes were used to perform a voting count for every potentially interacting pairs of residues between the two proteins. The vote increases by one every time a pair of residues *i* ∈ [1, *L_A_*] and *j* ∈ [1, *L_B_*] are within a threshold distance *d* Å from each other in a candidate complex as predicted by ClusPro. We experimented with different values of *d* from 8 to 12 *Å*. Since docking is an approximation, we settled for the cutoff of 12*Å* to get a larger pool of candidates for binding. The information obtained from this matrix is too sparse. However, it can roughly identify the neighborhood of binding sites. We utilized a smoothing techniques based on a Gaussian convolution filter with a kernel of size 3 to diffuse the information to neighboring residues. The smooth matrix is then turned into a probability *M* ^1^ ∈ *R*^*L*_*A*_×*L*_*B*_^ by normalizing the values.

### Propensity Scores

To construct a contact propensity map, we utilized a subset of the data for training (total of 58 pairs of interacting proteins [12]) and measured the frequency of contact between amino acid types. Contacting amino acids were defined as residue pairs with distance *<* 12Å, which is an upper bound for the default contact distance. We normalized the frequencies into a probability matrix in a similar fashion: *M* ^2^ ∈ *R*^*L*_*A*_×*L*_*B*_^. This matrix encodes the contact propensities between residues in our training set of proteins. The rationale behind this is that not all amino acids have an equal probability of interacting with other amino acids, and hence a propensity map provides prior information on potential of residues to interact. Figure 2 shows a heatmap of the propensity score.

**Figure 2.**
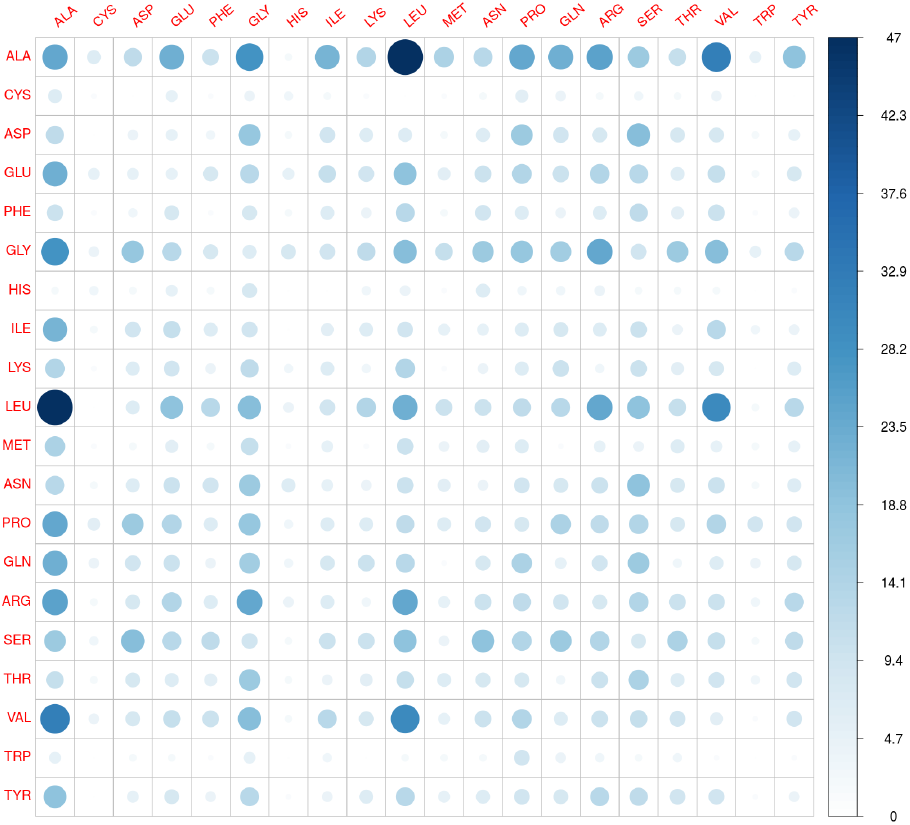
The amino acid contact frequency measured from all the training data. As can be seen the propensity of amino acids pairs can vary based on the type of amino acids. This data is converted into probability of interaction based on amino acid type. Two amino acids are in contact if the distance between their C-*α* < 12*Å*.

### Prior Information as a Penalty Matrix

The geometric and propensity contact maps were combined into a single penalty matrix as a weighted average *M* = *w*_1_*M* ^1^ + *w*_2_*M* ^2^*w*_1_ + *w*_2_. Here 0 ≤ *w*_*i*_ ≤ 1 denote the relative weights of the corresponding contact maps *M ^i^*, *i* = 1, 2. The weights reflect the significance of each contact map and are set as the accuracy of the contact map in predicting correct binding interfaces between protein complexes in the training set. The information in the probability model *M* is imposed as an *L*_1_ penalty in estimating the precision matrix Θ in a way that pairs with a high probability of contacting are penalized less. Note that even though we only used two sources of information in this work, the penalty matrix can readily be extended using more sources of information. This is the subject of current and future work. This information complements the information contained in the co-evolution data from the MSA, particularly for those residues with low co-evolution scores. Converting the probability matrix *M* into a correct form of penalty for estimating a sparse covariance matrix is a challenging task and has a high impact on the performance of the model and the connectivity of the underlying graph. To find the optimal value of the tuning parameter, we first determined an upper bound for Λ, denoted by *λ*_*max*_, defined as the minimum value for which the precision matrix is 0. This was carried out by inspecting the soft-threshold function in the coordinate descent algorithm [8] that is used for fitting the optimization problem 1. Setting *λ*_*max*_ = *max*(*abs*(*S*)), where *S* is the empirical covariance matrix, will achieve this task, resulting in a completely sparse fit. We showed this previously [11]. Once the upper bound is determined, the Λ matrix is constructed as follows:

- Λ_*j,i*_= Λ_*i,j*_ = *λ_max_*, where *i, j* belong to only one protein
- 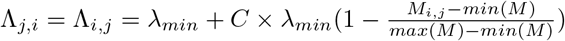, where, *i* and *j* belong to protein A and B, respectively or vice versa.

The value of *λ*_*min*_ is set to the threshold of convergence in the GLASSO, which is set to 0.0001. *C* is a positive integer, which was calculated using the training set and is set to 30. The elements of *M* are designed to force the values close to 0 if the probability of interaction between corresponding residues is high. For pairs of residues with low prior probability of interaction, the value is held near *C* × *λ*_*min*_. For all values of intra-protein residues, the corresponding values in *M* are set to *λ*_*max*_.

The matrix Λ is passed along with *S* to PSICOV. PSICOV, like other direct coupling methods, normalizes the precision matrix Θ by taking the norm 2 and applying an Average Product Correction (APC) [3] to remove phylogenetic biases. It then fits a logistic curve and ranks the top pairs. A caveat of setting the penalty for intra-residue pairs to *λ*_*max*_ is that the co-evolution score of corresponding residues within the single proteins is not taken into account, which in turn may impact the overall value of the normalized co-evolution during the calculation of APC. To address this issue, we took a “Meta” approach and ran PSICOV (and GREMLIN) without the penalty first, and filtered out the intra-protein pairs. We refer to these combined methods as **S-PSICOV** and **S-GREMLIN** respectively. This is similar to the approach taken in Meta-PSICOV [14], where combining the result of multiple methods (PSICOV, plmDCA, and GREMLIN) improves the performance of proteins folding.

### Post Processing by Patch Building and Filtering

As discussed in the Introduction, building interface patches of residues around co-evolving residues that are predicted to be at the interface should help in improving model performance and reducing false positive rate. In this section, we describe our proposed patch-building and filtering process that accurately identifies clusters of interacting pairs. As will be shown later, this method has a significant impact on decreasing the false positive rate (see Results).

Let Γ be the list of top *L* co-evolving pairs of residues and let *P*_*i*_ denote a patch of neighboring residues of the *i*-th residue in the protein (*A* or *B*) determined from the structure of the protein (e.g a PDB file). Each residue in *P*_*i*_ has a distance of less than 4Å from *i* and has an absolute SASA that is greater than 1 Å.

These patches are constructed for each residue in each protein independently. To each residue pair *e* = (*i, j*) in the list *G*, we assign two patches, one from protein *A*, denoted by *P*_*e*_ = *P*_*i*_, and from protein *B*, denoted by 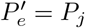. Next we construct a symmetric matrix *M*_Γ_ of size *L × L_i_*as follows:

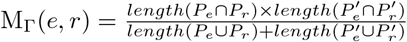, where *e* and *r* are residues pairs that appear in the list Γ. The elements of the matrix *M*_Γ_ are a modification of the Jaccard distance and are designed to measure the similarity between two patch-pairs 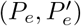 and 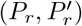. Note that the value *M*_Γ_(*e, r*) is non-zero if and only if there is an overlap between the two patches of residues *e* and *r* in protein *A* and also between the two patches of residues *e* and *r* in protein *B*.

The rational for this design is to distinguish between local clusters of interface pairs that have a relatively high co-evolution score and appear as neighboring patches of residues, and pairs of residues that are isolated nodes and have a high co-evolution score for other reasons such as MSA or phylogenetic biases. It is important to note that the binding site usually appears as a clusters of residues not as a single residue [**?**]. Once *M*_Γ_ is constructed, we remove the columns (rows) of *M*_Γ_ that consist of entirely 0s to filter top co-evolving pairs that have no intersection with the rest of pairs in Γ. This process will eliminate most of the false positive pairs from the final interface list, albeit a small fraction of true positive may also be filtered out.

Figure 3 shows the patch-building result (without removing any columns). *M*_Γ_ is represented as an adjacency matrix by replacing the non-zero elements with 1 and visualizing the resulting graph. Each pair is represented by a node in the graph. If the distance between two residues in a node is less than 12Å, it is colored in green and otherwise in purple. This graph shows that not all co-evolving residues correspond to physically interacting residues.

**Figure 3.**
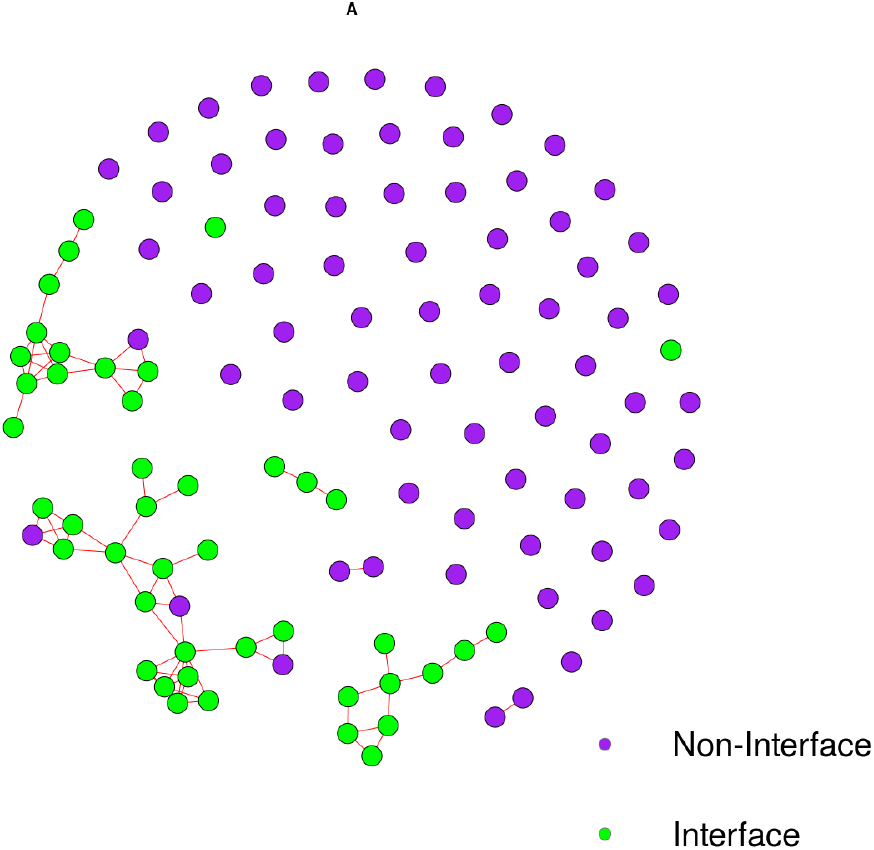
The patch matrix *M*_Γ_ is converted into an adjacency matrix for the top co-evolving pairs. The neighboring patches for 1RM6 complex between two chains B-C are shown. Green pairs correspond to actual interface residues (distance less than 12Å) and purple pairs are non-interface residues.

## Results

### Simulated Data

To test the general validity of our approach, we simulated a controlled dataset of MSAs with various degrees of co-evolution as follows: A first order HMM was used to generate an *in silico* MSA. To set the background probabilities, we used the PAM and BLOSUM62 substitution matrices and estimated the distribution of amino acids in all the PDB structures. Using this procedure, we built an MSA of size 1000 *×* 200. Next, six pairs of columns in the MSA were selected to model co-evolution. The strength of the co-evolution was tuned using three parameters as follows:

- Co-evolution parameter *α*: A score between 0 and 1, controlling the transition probability of a 21^2^ states HMM, with 0 corresponding to no co-evolution and 1 corresponding to maximum co-evolution.
- Conservation parameter: The rate of amino acid change from one type to another. 0 means that we expect to see no conservation, and 1 represents that co-evolution occurs between 2 amino acid types.
- Bias control: We have a fair bias which represents an original PAM matrix.

The six co-evolving pairs of columns were tuned using these parameters to generate multiple MSAs of various co-evolution strength going from low signal data (*α* = 0.05) up to a relatively strong signal data (*α* = 0.95). We then converted the MSA into a binary matrix as explained before. In our simulations, we used the maximum value of the covariance to set the max *λ* value. We then constructed a penalty matrix by setting the value of non-interacting pairs to *λ*_max_ while the rest of the columns received a value between 0.0001*λ*_max_ to 0.5*λ*_max_. The empirical covariance and the penalty matrix were then used to fit a GGM model. In our implementation, we utilized the GLASSO method implemented in

### Real Data

We trained and tested our method on benchmark datasets that were provided in the EV-Complex and GREMLIN [12, 26]. Each of the MSAs constructed for the datasets has at least one pair of proteins whose individual and complex protein structures are known, hence providing a ideal benchmark for our method. In each MSA, the first sequence was form *E. coli* and hence we used the *E.coli* complex as our input to the interface detection algorithm.

### Model Assessment

Protein-Protein interface prediction is an example of highly imbalanced problems, where the ratio of the positive class to the negative class is extremely small. Therefore, as in other approaches, we measured the precision 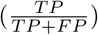 and 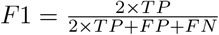 scores for all the models across different settings. Two ranking lengths for the assessing of top-ranked pairs were considered as follows:

- **Top-8**: This represents the total number of residues between two proteins in a complex that have a distance of less than 8Å from each other.
- **Top-12**: This represents the total number of residues between two proteins in a complex that have a distance of less than 12Å from each other.

We also considered 8 and 12Å as two different interaction cutoffs. In order to evaluate the impact of structural prior on DCA models, measuring the relative metric with respect to existing methods is recommended where the relative F1 and precision scores are:

- 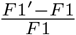
- 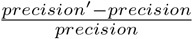

Positive and negative values of the relative scores correspond to gain or loss of improvement of the new model as compared with the base model. We benchmarked five different models to assess contribution of the penalty matrix and patch-building on model performance. The first two are Meta-methods that incorporate structural-based prior penalty with PSICOV and GREMLIN which are denoted as **S-PSICOV** and **S-REMLIN**. The next two models incorporate the patch-building and filtering process in addition to the structural information and are denoted by **PatchS-PSICOV** and **PatchS-GREMLIN**. Finally, to investigate the impact of patch-building and filtering independent of the structural prior, used the patch process subsequent to GREMLIN without any structural information. This method is denoted by **Patch-GREMLIN**. We did not include Patch-PSICOV in the result section since it is known that the performance of DCA in Pseudo-likelihood is greater than the GGM [25] (Our benchmark also confirmed this). The performance of the models were assessed on 19 complexes. We did not compared any deep learning models such as RaptorX as these models use a training set that include several of the sequences in our training and test sets. Large training sets of few thousand protein MSAs are required to train any deep learning models, which make re-training and testing on benchmark data very difficult. Therefore, one of the advantages of our method is that it does not rely on a large set of training unlike deep learning approaches. Figure 4 shows the average relative F1 score of all five proposed models with respect to PSICOV (black) and GREMLIN (gray). In Figure 4 (A) and (B), the binding site cutoff is set to 8 and 12Å and the length of top co-evolving residues for measuring F1 score for each complex is based on the Top-8 and the Top-12 respectively. As can be seen PatchS-PSICOV improved the F1 score relative to PSICOV and GREMLIN by 70% and 24% respectively. Although the structure-based prior models do not show much improvement by themselves, after the patch-building and filtering the relative improvement increases significantly. All the patch-based models results in improved relative F1 performance. This demonstrates the impact of integrating structural prior along with patch-filtering (compared to Patch-GREMLIN, which does not utilize structural information). This is because the prior information by itself adds a lot of false positive and true positive pairs to the top co-evolving pairs but when patch-building is applied, those pairs that are isolated (mostly false positive) will be eliminated from final list. In a similar fashion, we measured the relative precision for all five new models with respect to both PSICOV and GREMLIN. Figure 5 shows the average relative precision score. As in the F1 score, all patch-based models improve the average relative precision. Supplementary Figure 2 compares the top 100 ranked pairs against the actual distance in E.coli F1-ATP synthase inhibited by subunit Epsilon complex (PDB 3OAA), chains G and H. As shown, there exist more residue pairs within binding site distance in all the new proposed models relative to the base model. This is an example of how structural information and patch refinement can impact the performance of a model (contact distance is assumed 12Å).

**Figure 4.**
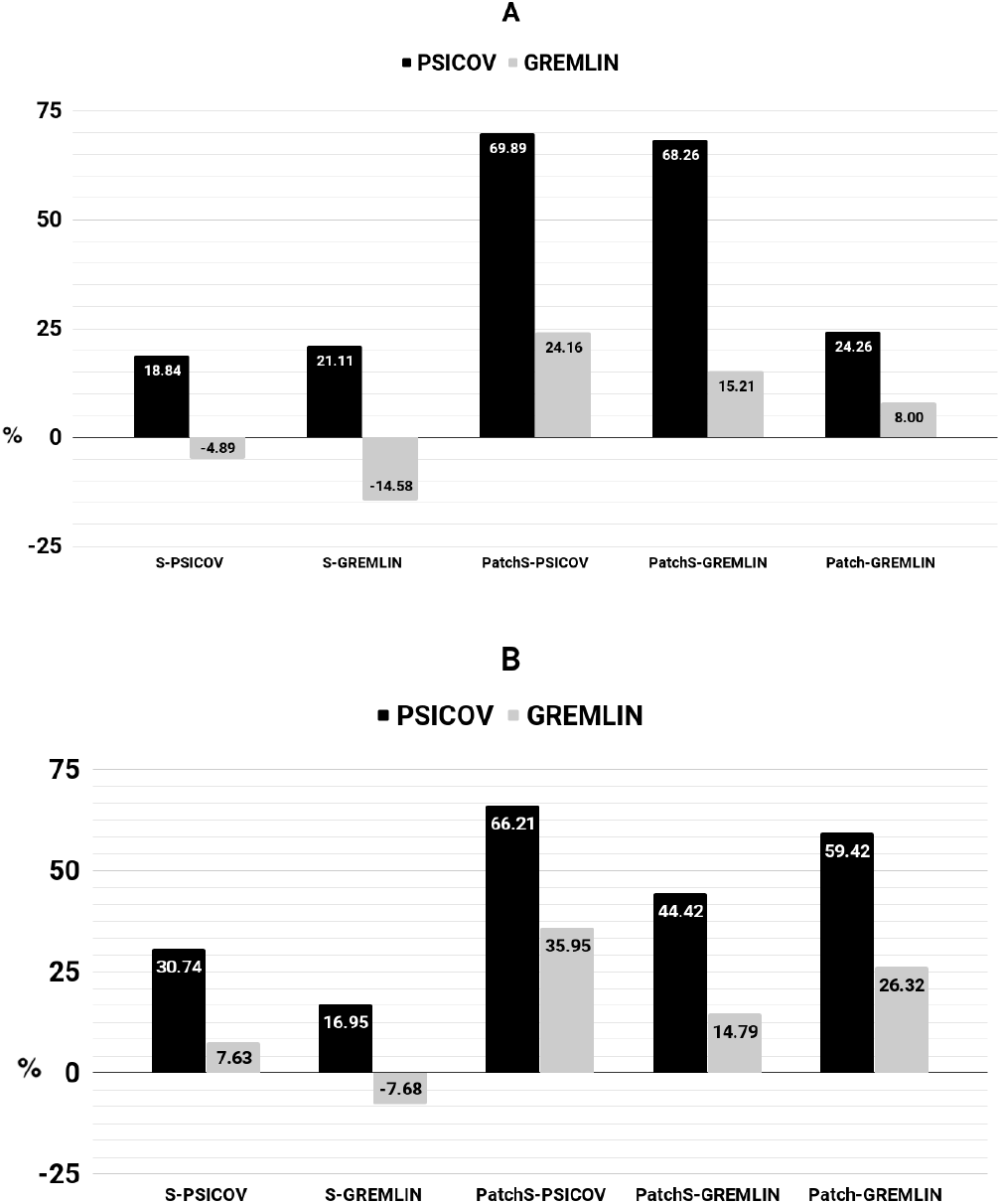
Average relative F1 score of the new proposed models compared to PSICOV and GREMLIN on the 19 protein complexes. The binding site threshold is set to 8Å (A) and 12Å(B) respectively.

**Figure 5.**
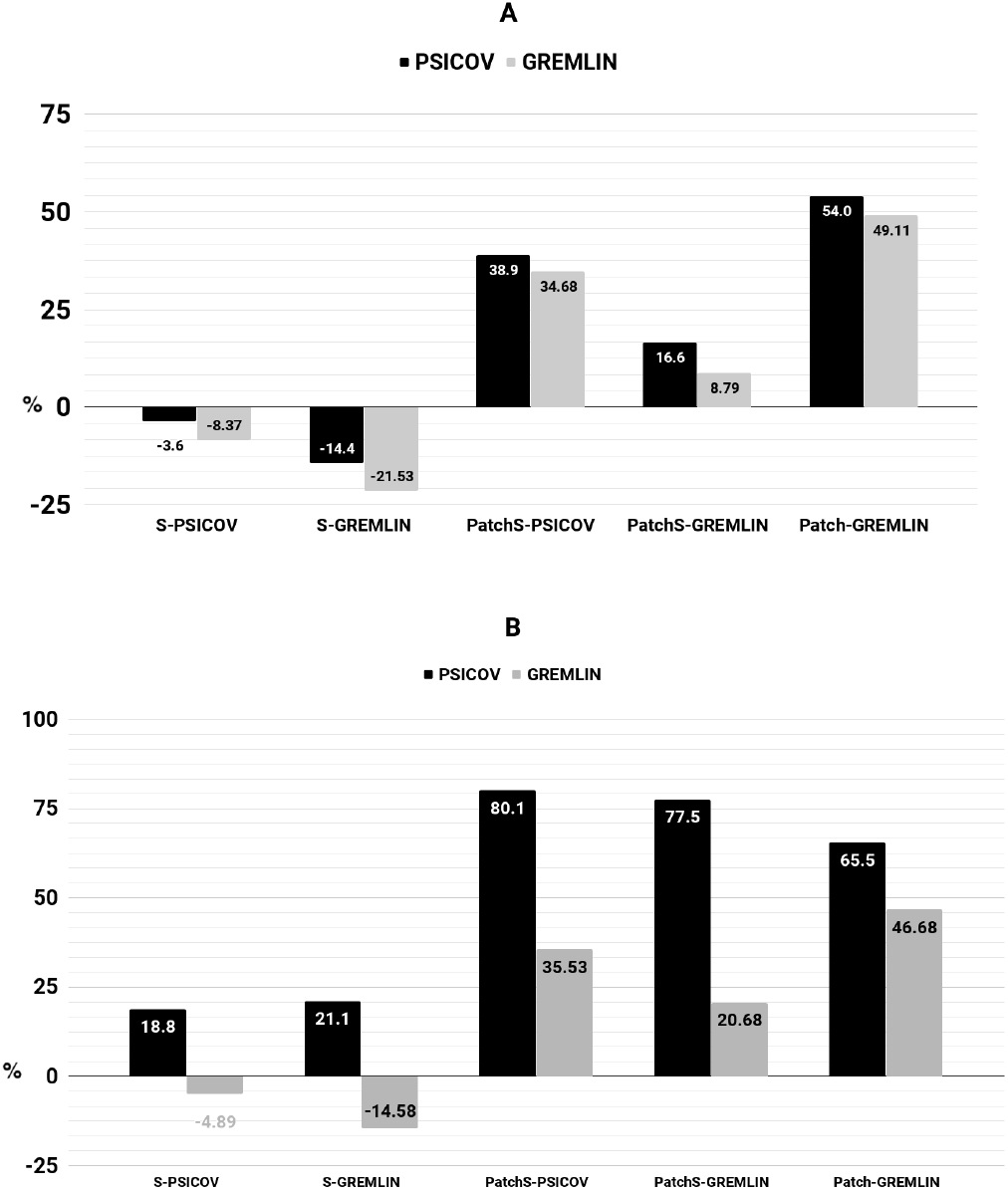
Average relative precision score comparison of the new proposed models with PSICOV and GREMLIN. In A and B the binding site threshold are set to 8Å and 12Å, respectively among top Top-8 pairs.

Table 1 shows the average F1 score for all proposed models as well as PSICOV and GREMLIN. As can be seen, the average F1 score in Patch-GREMLIN outperforms all other models under all three different settings. Similarly, table 2 represents the average precision scores. The Patch-GREMLIN model outperforms all other models under two different conditions. In both conditions there is significant improvement in average precision score compared to other methods. Moreover, PatchS-PSICOV outperforms PSICOV and GREMLIN models for binding site cutoff 12Å. We also evaluated the performance of geometry-based model (the *M* ^1^ matrix) calculated from ClusPro to examine the influence of the docking on the performance of our models. As can be seen, the performance of geometry-based model is relatively low on its own, but when this information is integrated into the PSICOV followed by patch-building, the performance improves significantly.

**Table 1.** Average F1 score comparison between different models under different conditions

| Model\Condition | cutoff=8,<br>Top-8 | cutoff=12,<br>Top-8 | cutoff=12,<br>Top-12 |
| --- | --- | --- | --- |
| GREMLIN | 0.20 | 0.43 | 0.16 |
| PSICOV | 0.18 | 0.39 | 0.13 |
| S-PSICOV | 0.13 | 0.32 | 0.15 |
| S-GREMLIN | 0.10 | 0.27 | 0.13 |
| PatchS-PSICOV | 0.17 | 0.41 | 0.18 |
| PatchS-GREMLIN | 0.14 | 0.34 | 0.16 |
| Patch-GREMLIN | **0.24** | **0.48** | **0.20** |

**Table 2.** Average Precision score comparison between different models under different conditions

| Model\Condition | cutoff=8, Top-8 | cutoff=12, Top-12 |
| --- | --- | --- |
| GREMLIN | 0.20 | 0.43 |
| PSICOV | 0.18 | 0.39 |
| geometry-based model | 0.17 | 0.31 |
| S-PSICOV | 0.13 | 0.32 |
| S-GREMLIN | 0.10 | 0.27 |
| PatchS-PSICOV | 0.19 | 0.45 |
| PatchS-GREMLIN | 0.15 | 0.36 |
| Patch-GREMLIN | **0.32** | **0.62** |

## Discussion

Taken together, our results indicate that incorporating structural information can improve the performance of DCA-based models for predicting binding site between two interacting proteins. This is especially important in larger proteins where the search space increases exponentially. The structural information used in our method is based on an ensemble combination of protein docking and propensity, which are general properties for all protein families and it can be expanded to incorporate more models. The docking model is based on geometric complementarity of partner proteins and energy function. The binding characteristics for each pair of residues obtained from proteins structures is used as prior knowledge and incorporated into a co-evolution GGM model as a penalized optimization. Further, our patch-based filtering approach results in significant improvement of model performance and elimination of many false positives. As a future work, we plan to utilize comprehensive algorithms to further probe and interrogate the structure for additional information. For example by integration more features such as secondary structure and rigidity analysis of the protein structures, we expect to improve the overall performance of the model. Such improvements can also be utilized in deep learning models to obtain a more precise models.

## Supporting Information

## Acknowledgments

We would like to thank Sergey Ovchinnikov and Michael Tolstorukov for their help and guidance for this project.

