## Supplemental File for "Patch-DCA: Improved Protein Interface Prediction by utilizing Structural Information and Clustering DCA scores"

**Supplementary Material 1**

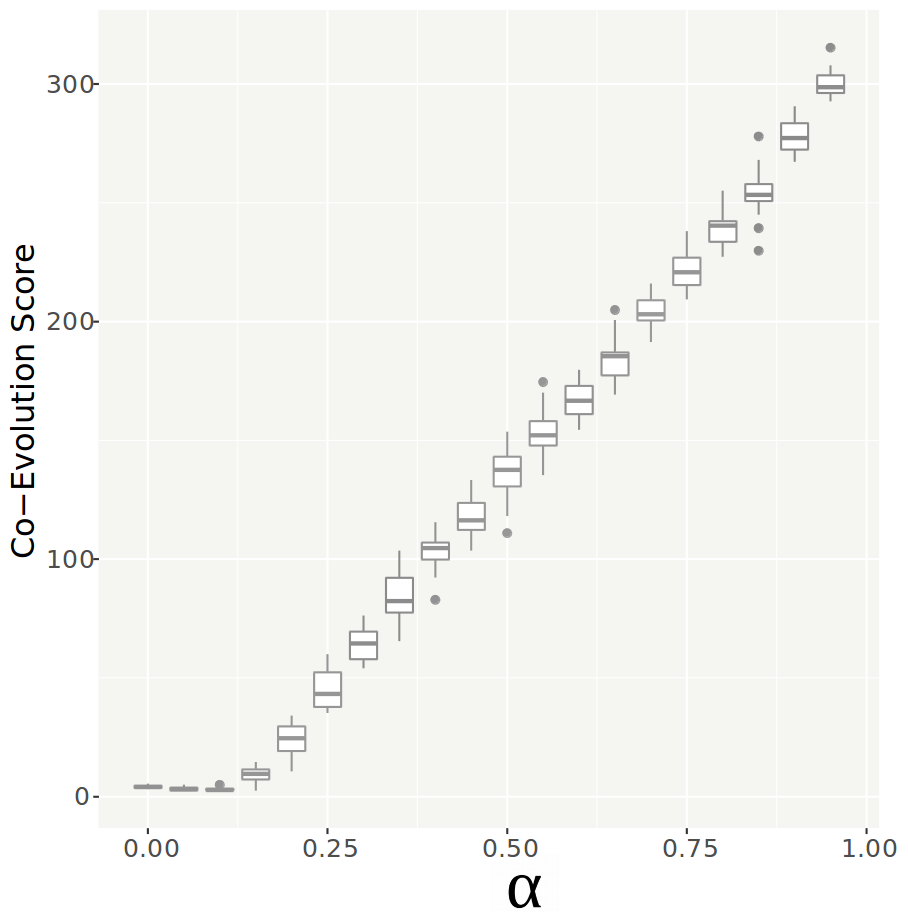

**Supplementary Fig1:** The GLASSO score as a function of α, the co-evolution parameter for interacting

pairs. The error bars are shown near the boxes. y-axis represents an unnormalized GLASSO

score obtained from each complex.

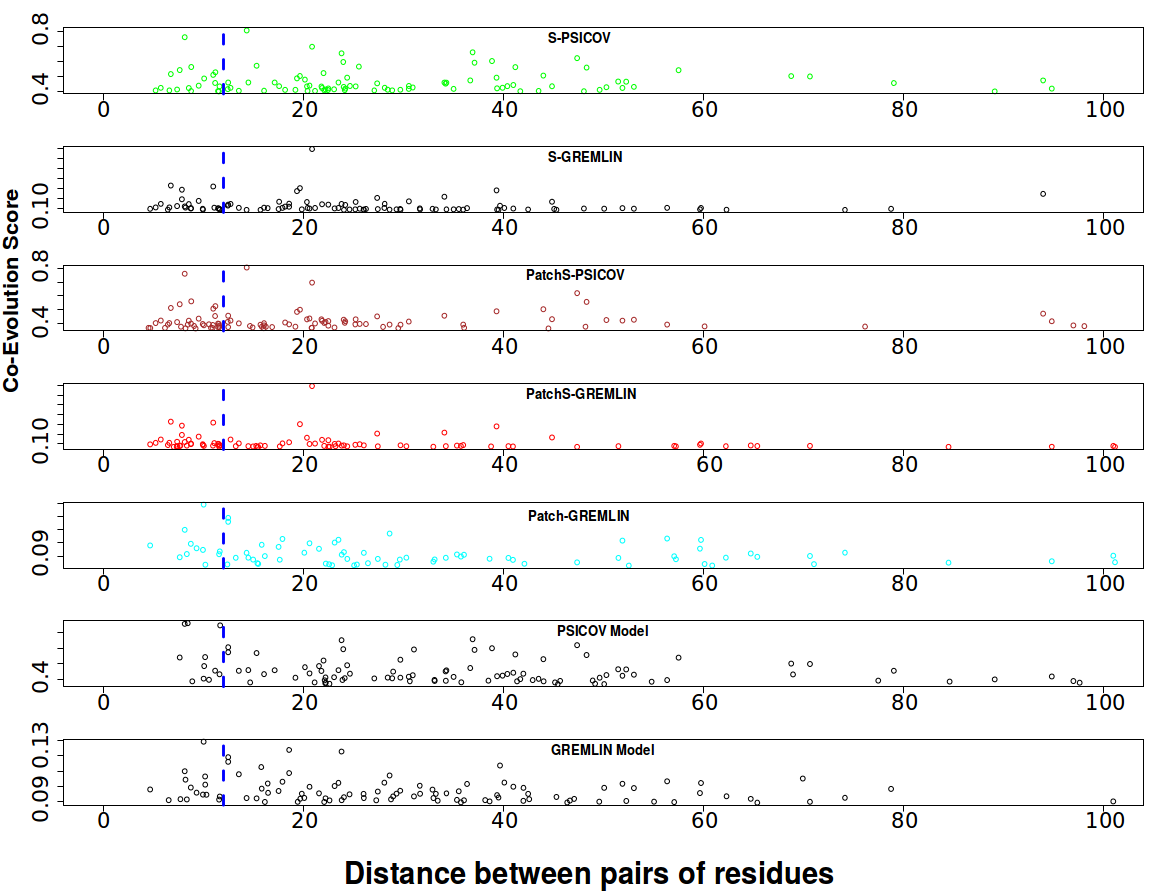

**Supplementary Fig 2**: The relationship between co-evolution score (y-axis) in top 100 contact pairs with respect to the distance (x-axis) between two residues in the 3OAA complex among all the models. Blue line represents the cutoff which is 12 Å.

Supplementary Table 1: List of the test set complexes

| **Protein (pdb code)** | **#of residues in Chain A** | **#of residues in Chain B** | **Number of Residues within 8 Angestrom (Top-8)** | **Number of Residues within 12 Angestrom (Top-12)** |
| --- | --- | --- | --- | --- |
| 1B70_A_1B70_B | 265 | 776 | 189 | 1095 |
| 1EFP_A_1EFP_B | 307 | 247 | 77 | 430 |
| 1EP3_A_1EP3_B | 311 | 262 | 39 | 253 |
| 1QOP_A_1QOP_B | 265 | 391 | 42 | 308 |
| 1RM6_A_1RM6_C | 761 | 158 | 48 | 304 |
| 1RM6_B_1RM6_C | 323 | 158 | 46 | 273 |
| 1TYG_B_1TYG_A | 65 | 243 | 26 | 245 |
| 2NU9_A_2NU9_B | 285 | 386 | 68 | 531 |
| 2ONK_A_2ONK_C | 240 | 253 | 43 | 309 |
| 2VPZ_A_2VPZ_B | 734 | 194 | 16 | 105 |
| 2WDQ_C_2WDQ_D | 121 | 106 | 15 | 175 |
| 2Y69_A_2Y69_B | 513 | 228 | 24 | 163 |
| 2Y69_A_2Y69_C | 513 | 260 | 16 | 57 |
| 3IP4_A_3IP4_B | 485 | 483 | 11 | 55 |
| 3IP4_B_3IP4_C | 482 | 93 | 62 | 278 |
| 3MML_A_3MML_B | 285 | 208 | 32 | 175 |
| 3OAA_H_3OAA_G | 138 | 285 | 105 | 623 |
| 3PNL_A_3PNL_B | 356 | 212 | 22 | 174 |
| 3RRL_A_3RRL_B | 227 | 198 | 47 | 271 |

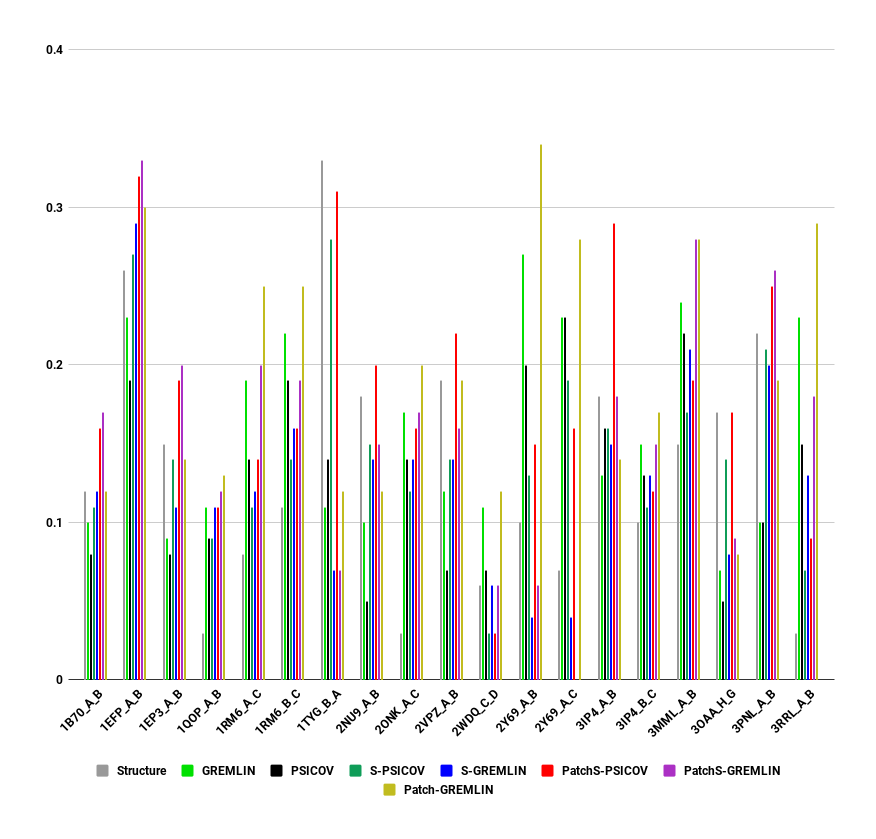

**Supplementary Figure 3:** The comparison of F1 score across test set complexes. The binding site distance set to 12 A and the top B-rank pairs are selected.
